# Genome Sequencing of Rice Landraces from the Yuanyang Terraces Uncovers Ancient and Diverse Lineages of Indica Rice

**DOI:** 10.1101/2025.09.09.675106

**Authors:** Huichuan Huang, Shengchang Duan, Xiang Li, Linna Ma, Xiahong He, Youyong Zhu, Chengyun Li, Yang Dong, Jean-Benoit Morel, Pierre Gladieux

**Affiliations:** State Key Laboratory for Conservation and Utilization of Bio-Resources in Yunnan, Yunnan Agricultural University, Kunming, China; Key Laboratory for Agro-Biodiversity and Pest Control of Ministry of Education, Yunnan Agricultural University, Kunming, China; Yunnan Research Institute for Local Plateau Agriculture and Industry, Kunming, China; Key Laboratory of Forest Resources Conservation and Utilization in the Southwest Mountains of China Ministry of Education, Southwest Forestry University, Kunming, China; PHIM Plant Health Institute Montpellier, University of Montpellier, INRAE, CIRAD, IRD, Institut Agro, Montpellier, France

**Keywords:** domestication, landraces, rice, population genomics, varietal mixing

## Abstract

High-yielding elite rice cultivars exhibit limited genetic variability, raising concerns about our capacity to sustain productivity in the face of changing biotic and abiotic threats. Meeting the challenges that lie ahead largely depends on our ability to make use of novel sources of genetic variation and re-engineer agrosystems. Here, we report on the evolutionary history and population genetic structure of 353 accessions representing 91 landraces from China’s centuries-old Yuanyang terraces of rice paddies (YYT). We found that the indica YYT landrace population is genetically structured and exhibits high standard variation. Analysis of natural selection reveals that innate immunity genes have a marked difference in coevolutionary dynamics between modern and traditional rice, characterized by a stronger influence of directional selection, which reduces diversity, in modern varieties. Our study highlights the importance of preserving landraces and the need for targeted efforts to integrate the standing variation in landraces into new varieties.

## Introduction

Asian rice landraces, *i*.*e*., rice populations that have experienced sustained cultivation in traditional agrosystems, are thought to represent genetic intermediates between wild *O. rufipogon* progenitors and modern elite *O. sativa* cultivars (*1*). Having been selected for genetic variants and gene combinations that are favorable in a wide range of geographic localities and cultural contexts, rice landraces are expected to represent a rich reservoir of genetic variation, readily accessible to breed new varieties with superior agronomic performance (*2*). Recent population genomic surveys of large collections of wild accessions, modern varieties, and landraces have contributed to clarifying the genetic makeup and population structure of *O. rufipogon* and *O. sativa* (*3-6*), and refining the domestication and migration history of japonica and indica landraces (*7*). However, despite the agronomic and academic interest in landraces, our understanding of the dynamics of rice genetic diversity on a local scale and the impact of the Green Revolution on such local varietal landscape has remained limited (*8, 9*). The emblematic Yuanyang terraces (YYT) of Yunnan (China) represent an outstanding experimental set-up to investigate the factors underlying the maintenance of genetic diversity in crops and its relationships with local cultures (*10, 11*). The Yuanyang terraces cover ∼21,000 ha and harbor more than 195 rice landraces (*12*) that have been grown for more than 1300 years (*13, 14*), while relatively little affected by infectious diseases (*15, 16*).

## Results and Discussion

We generated and analyzed whole genome data for 353 accessions, representing individual plants of the Yuanyang terraces randomly selected in 91 bags of seeds with different varietal names given by farmers. This dataset was supplemented with genomic data for a subset of 437 accessions from the 3,000 rice genomes project (*4*), as well as 42 additional varieties representing different rice sub-species (collectively hereafter referred to as ‘modern’; Dataset S1). The full dataset was composed of 832 genomes and sequenced at an average depth of 14.5× for the YYT dataset. All sequences were aligned against the *Oryza sativa* indica genome R498 (*17*), and SNP calling identified 17.3 million polymorphic sites after filtering.

Genome-wide variability was highly structured in our dataset. Principal-component analysis (PCA) partitioned the accessions into three clearly defined groups. The first principal component (7.5% of variation) separated japonica and indica varieties (Figure 1B). Several accessions, most of which were sampled in the YYT, were genetically intermediate between indica and japonica, and may represent hybrids between the two rice subspecies. The second principal component (5.5% of variation) separated 3KGP and modern varieties from YYT landraces, indicating that the Yuanyang rice landraces represent a large and original reservoir of genetic diversity (Figure 1B).

**Figure 1.**
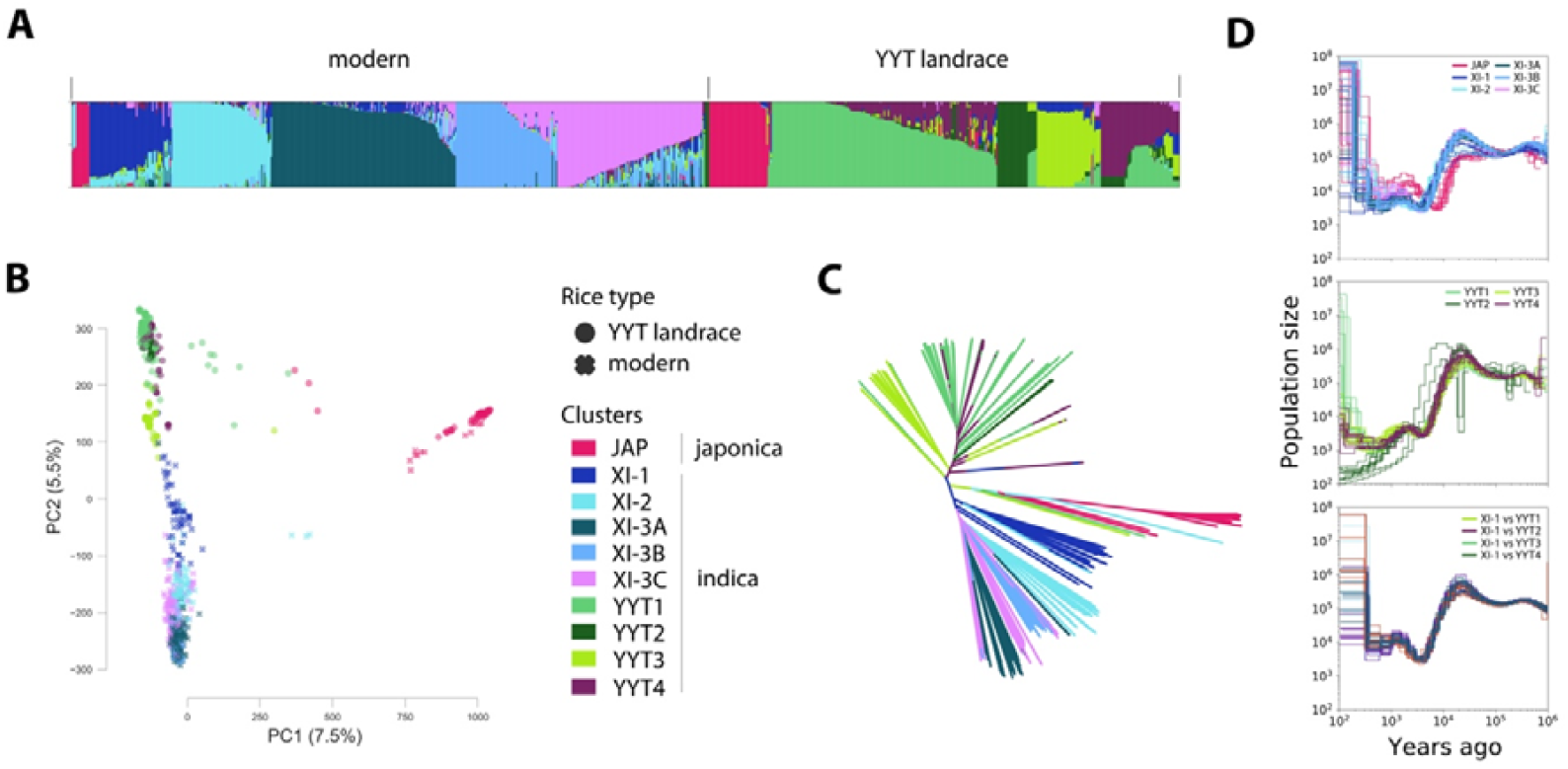
Population structure and population dynamics of 353 landrace accessions from the Yuanyang terraces and 479 accessions of modern Asian rice. **(A)** Stacked bar plot representing the percentage of membership (q) for each cluster identified at K = 10. The color designations for each cluster illustrated in **(A)** are used to represent accessions in all the other panels (**B-D**). An accession was assigned the color of the cluster of the highest q value. Cluster designations are shown in panel **B. (B)**, Principal component analysis of 4.6e6 single nucleotide polymorphisms in 832 accessions. The plot shows the first two principal components and the percentage of explained variation. **(C)** Neighbor-joining dendrogram based on p-distance between 832 accessions (p-distance is the number of differences divided by the number of observations). **(D)** Effective population size as a function of time, as estimated using the coalescent approach implemented in MSMC2 (*19*), with a mutation rate of 6.5e-9/bp/generation (*20*) and a generation time of one year. The scale for both the x and y axes is logarithmic (base 10). Modern indica and japonica accessions are shown in the top subplot, indica landraces in the middle subplot, and cross-cluster analyses in the bottom subplot. In single-cluster analyses (top and middle subplots), each line represents a set of eight haplotypes, each haplotype representing a different accession. In cross-cluster analyses (bottom subplot), each line represents a set of eight haplotypes, with four haplotypes from each cluster, each haplotype representing a different accession.

We explored the major groups defined by PCA using model-based clustering (*18*) with Admixture. Cross-validation monotonously decreased with increasing K and thus did not point to a particular number of clusters as modeling the data more adequately (Figure S1). However, coefficients of ancestry revealed that K=10 clusters was the smallest K value that (i) showed an alignment of membership patterns with accession labels (i.e., 3KGP and modern indica were distinct from indica accessions from Yuanyang terraces), (ii) clearly individualized the previously identified clusters XI-1, XI-2 and XI-3 (*4*) in the modern and 3KGP dataset, and (iii) identified two clusters of Yuanyang accessions known to result from recent crop improvement (YYT2 and YYT4). Further increasing K mostly distinguished smaller clusters within the major clusters identified at K=10 (Table S1; Figure S2). Among the K=10 clusters featured: (i) one Japonica cluster, containing both japonica from YYT and 3KGP, hereafter referred to as JAP; (ii) four clusters of Yuanyang landraces (collectively referred to as YYT clusters): YYT1, YYT2, YYT3 and YYT4; (iii) five clusters of modern and 3KGP indica varieties (collectively referred to as XI clusters): XI-1, XI-2, XI-3A, XI-3B and XI-3C, with the latter three clusters corresponding to the subdivision of cluster XI-3 (Figure 1A).

Indica landraces from the YYT formed a distinct group of indica rice. In the neighbor-joining dendrogram, japonica landraces from the YYT grouped with other japonica accessions, while YYT indica landraces formed a lineage distinct from other modern or 3KGP indica accessions (Figure 1C). The average nucleotide divergence between YYT clusters and XI clusters (from dxy=0.0012 to dxy=0.0028) was on average two-thirds the divergence between XI clusters and the japonica cluster (dxy=0.0026 to dxy=0.0037), which separated from 350 to 400,000 years ago (*21-23*) (Table S1). When tracing the demographic history using multiple sequentially Markovian coalescent approach (Figure 1D), YYT clusters showed a strong bottleneck at the same period (3000 to 6000 years ago) as the XI clusters, but younger than the bottleneck detected for japonica (7000 to 10000 years ago). These demographic events are likely associated with the initial domestication of indica rice, and they are consistent with japonica domestication being older than indica domestication (*7, 21, 24*). YYT1, YYT2 and YYT3 clusters also experienced an intense bottleneck and subsequent population growth between 500 and 1000 years before present, consistent with anthropological studies dating the foundation of the YYT agrosystem to 1300 years ago (*13*). Estimates of effective population pointed to a recent and intense bottleneck in cluster YYT2, confirming information provided by local farmers, who described the varieties in this cluster as recently released, improved landrace lines. Admixture analyses also support the relative isolation of YYT landraces from the remainder of indica rice diversity, as represented in our dataset by the XI clusters. While six of the 437 3KGP genomes had high membership proportions in YYT clusters (Dataset S2), indicating that Yuanyang landraces were represented in the 3KGP dataset, none of the YYT accessions had high membership in XI clusters. Only one YYT accession showed membership proportions consistent with recent admixture with XI clusters (accession 123; Dataset S2). Other signatures of possible admixture appeared older and indicated a history of breeding with non-YYT germplasm in the cluster YYT3 of improved landraces, as well as, to a more limited extent, in the YYT4 cluster (Figure 1C; Dataset S2). Together, these analyses show that indica YYT landraces form a divergent, early diverging lineage of indica rice, with a distinctive demographic history.

Coexistence and admixture among genotypes from different YYT clusters were common in bags of seeds. Among the 353 YYT accessions, 33% had mixed ancestry, with 111 accessions having ancestry in multiple YYT clusters, and five accessions identified as japonica × indica hybrids. Among the 69 samples of four seeds collected in bags of seeds with different names, 68% included at least one accession identified as a mixed ancestry genotype, and 39% included accessions from at least two different categories, with categories representing clusters, groups of admixed genotypes, or groups of hybrid genotypes (Figure 2). Deep sequencing of the most common landrace, *Acuce*, confirms the high lineage heterogeneity within bags of seeds and the high genomic diversity associated with a landrace name (*11*). These results indicate that genotype mixtures with paddy fields are a major component of the genetic heterogeneity of the YYT agrosystem.

**Figure 2.**
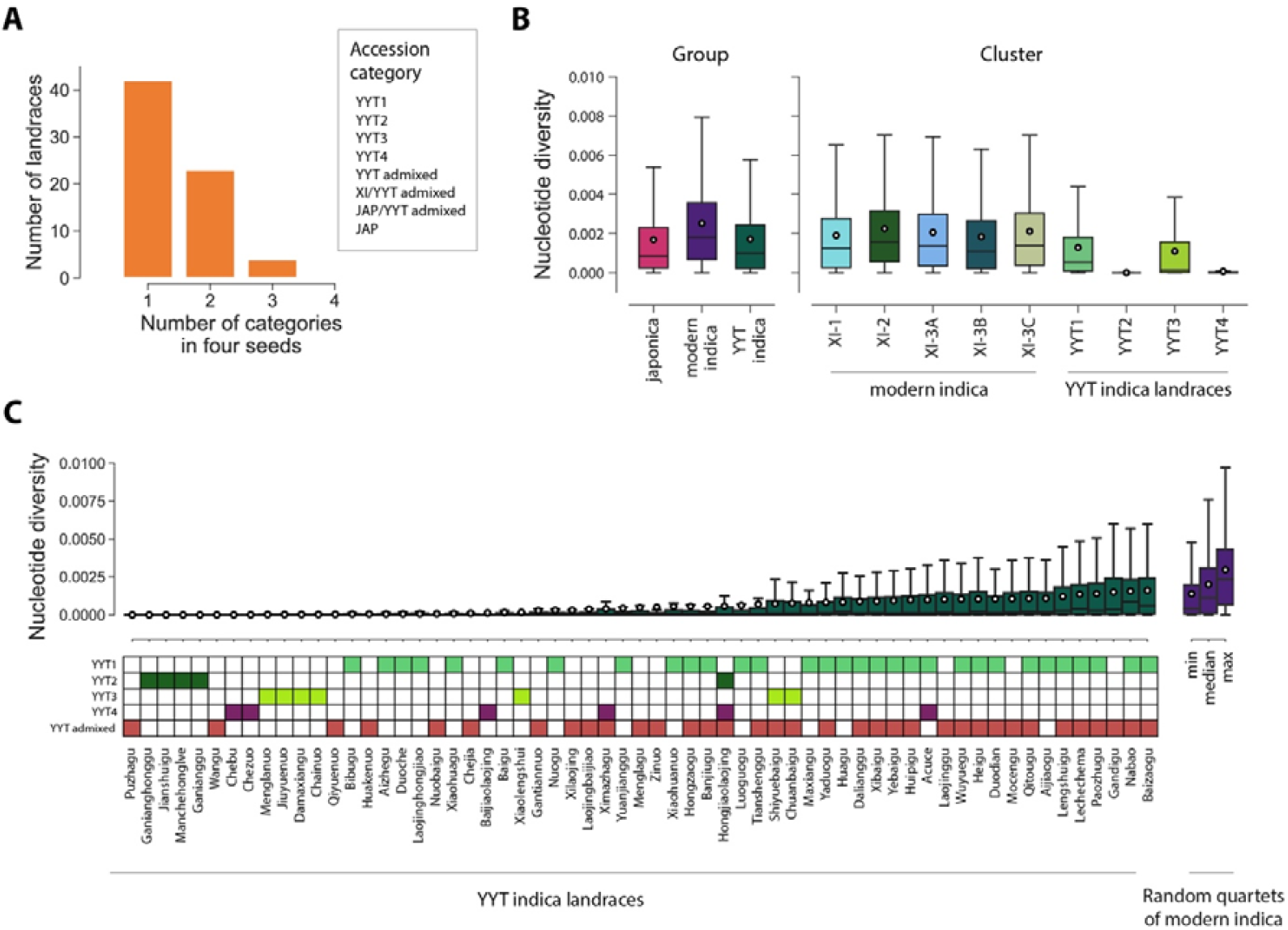
Nucleotide diversity in YYT landraces and modern varieties. **(A)** composition bags of seeds representing 69 landraces. Only landraces with a sample size of four were included in this analysis. Accessions were assigned to categories based on the membership proportions in the clustering analysis presented in Figure 1A. **(B)** nucleotide diversity in 10 kb windows in groups and clusters of accessions. All differences were statistically significant (Kruskal-Wallis tests followed by posthoc Mann-Whitney with Bonferroni-Holm correction, p<0.05). **(C)** nucleotide diversity in 10 kb windows in 60 individual landraces and 113 randomly sampled quadruplets of modern indica varieties. The names used to designate landraces were reported by farmers when bags of seeds were sampled. Each landrace represents a different bag of seeds. The average nucleotide diversity per landrace was calculated when at least four representative genomes sampled from the same bag of seeds were available. In **(C)**, japonica and japonica/indica hybrid varieties were excluded.

Despite an overall high nucleotide diversity at the scale of the agrosystem, the YYT terraces were very heterogeneous in terms of varietal diversity. The level of nucleotide diversity in indica landraces (π=0.00171/bp) was comparable to the level observed in a global collection of japonica rice (π=0.00168/bp), and represented 68% of the level observed in a global collection of indica varieties (π=0.00251/bp). At the cluster scale, clusters of traditional, non-improved landraces YYT1 (π=0.00131/bp) and YYT3 (π=0.00106/bp) harbored from 47% to 58% of the standing variation observed in the most diversified XI cluster (XI-2: π=0.00225/bp) (Figure 2B; Table S2). By contrast, clusters of recently improved landraces harbored very little standing variation (YYT2: π=1×10^-6^/bp; YYT4: π=8×10^-5^/bp). Consistent with the differences observed across clusters, we found that highly variable landraces (i.e., landraces whose nucleotide diversity is of the same order of magnitude as random quartets of 3KGP indica varieties) coexisted with landraces that appeared clonal (Figure 2C). Mixtures in seed bags appeared to contribute to this pattern, as seed bags with the greatest nucleotide diversity tended to contain genotypes from multiple clusters, or genotypes that resulted from admixture (Figure 2A,C). The varietal heterogeneity in diversity levels suggests differences in breeding system (i.e., morphological or physiological aspects of sexual compatibility (*25*)), which may induce differences in mating system (i.e., levels of selfing vs outcrossing). Our results also indicate that traditional agricultural practices have allowed the maintenance of high levels of standing genetic variation in the Yuanyang terraces compared to global collections of indica plants.

The level of variation maintained in landraces was particularly remarkable for immunity genes. In all groups and clusters, nucleotide diversity was consistently and significantly higher at NOD-like receptors (NLRs, of which R-genes represent a subset), wall-associated kinases (WAKs), and receptor-kinases (RLKs) than at other genes (Figure 3A; Table S3; Kruskal-Wallis tests followed by post-hoc Mann-Whitney U tests, p<0.05). The variation observed in indica landraces at immunity genes was higher than in a global collection of japonica plants and up to 82% (RLKs) of the variation observed in a global collection of modern indica plants (Figure 3A; Table S3). It further appears that the variation observed in landraces was unique and original, with 6.5% of the 3.79×10^4^ observed polymorphisms at immunity genes being exclusive to YYT indica, as well as the observation of exclusive haplotypes and high genetic diversity at R-genes and immune receptors (Figure 3A-C). Allele frequency spectra of shared polymorphisms at NLRs showed enrichment in intermediate frequency alleles, indicating that the shared polymorphisms are older alleles maintained by rare allele advantage and not recurrent mutations, which are expected to segregate at low frequency (Figure 3E). However, examining ß-score correlations between modern and landrace varieties shows that genes exhibiting signatures of strong balancing selection tend to differ between the two groups, which could reflect differences in pathogen communities in the two agrosystems (Figure 3F). Even if our whole-genome sequencing approach did not recover the full extent of nucleotide diversity that can be characterized with NLR sequence-capture data (*28*), our study, based on almost one-half of existing landraces in the YYT, shows that extensive genetic diversity concerns not only the NLRs but also the other important classes of receptors, the WAKs and the RLKs.

**Figure 3.**
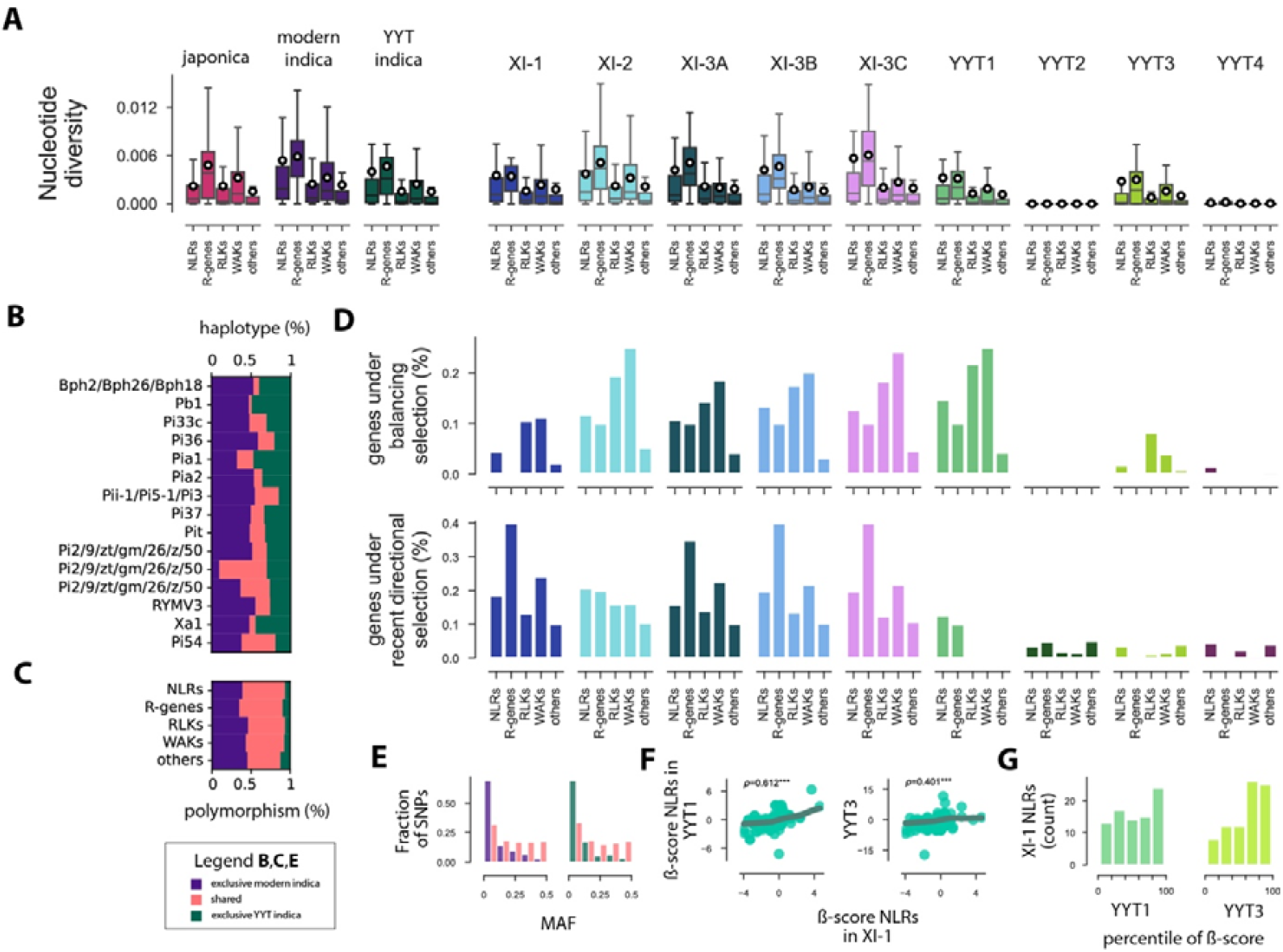
Standing variation and signatures of natural selection at immunity genes in modern Asian rice varieties and YYT landraces. **(A)** nucleotide diversity per base pair at NLRs, resistance genes (R-genes, which represent a subset of NLRs), RLKs, WAKs, and other genes in groups and clusters of modern varieties and YYT landraces. In box plots, circles represent the mean. **(B)** distribution of predicted protein haplotypes at R-genes. **(C)** proportions of shared and exclusive polymorphisms. **(D)** number of genes with signatures of balancing or recent directional selection. Genes under balancing selection are genes located in genomic windows in the top 5% of the ß-score, as estimated using Betascan 2 (*26*); genes under balancing selection are genes located in genomic windows of the top 5% of the XP-NSL statistic, as estimated using Selscan 2 (*27*). **(E)** allele frequency spectra at shared and exclusive SNPs. MAF is the minor allele frequency. **(F)** ß-score of NLRs in clusters YYT1 and YYT3 as a function of the ß-score of NLRs in cluster XI-1. ρ is Spearman’s rank correlation coefficient (***p<0.001). The curves represent nonparametric locally weighted linear regressions (lowess model). **(G)** Distribution of percentile of ß-score in clusters YYT1 and YYT2 for NLRs under recent directional selection in XI-1.

High polymorphism at immunity genes was not ubiquitous, however, with all groups and most clusters exhibiting a combination of weakly and highly polymorphic immune receptors, which indicates that the evolutionary dynamics of immunity genes is not simply one of cycling processes caused by the advantage of rare alleles (trench warfare model), but also implies recurrent selective sweeps (arms race model). Using genome-wide scans to detect selective sweeps and balancing selection, we were able to quantify the contribution of the two types of selection in shaping immune receptor diversity. Genomic regions under balancing selection were significantly enriched in NLRs, WAKs, and RLKs in all clusters of modern indica and cluster YYT1 (Table S4; Dataset S3; Fisher’s exact test, p<0.0001), but the number of immunity genes under balancing selection was higher in YYT1 (153 of 850 genes) than in modern indica (min: 61 genes, cluster XI-1; max: 134 genes, clusters XI-2 and XI-3C; Table S4; Dataset S3; Figure 3D). By contrast, the number of immunity genes under recent directional selection was much higher in modern indica (min: 135 genes, cluster XI-3A) than in YYT indica (max: 81 genes; cluster YYT1; Figure 3D; Table S5; Dataset S4). Among NLR genes exhibiting signatures of directional selection in modern varieties, a substantial fraction exhibited signatures of balanced selection in landraces (Figure 3G). Differences in selection regime at immunity-related genes between modern varieties and landraces may arise through differences in pathogen eco-evolutionary dynamics, related to breeding and farming practices. Elite rice varieties result from intense selection aimed at improving yield and quality under relatively low-stress conditions, and modern agrosystems are relatively static in varietal composition and consist of vast areas of low immune diversity. By contrast, Yuanyang indica landraces have been selected for their capacity to provide stable yields in specific environmental conditions and under low-input agriculture, and landraces circulate widely with the Yuanyang agrosystem through the practice of seed exchange between farmers (*29*), which contributes to maintaining high immune diversity on a relatively small surface area.

Previous studies on rice blast, the major fungal pathogen of rice, have shown that in the traditional YYT agrosystem (*30*), and unlike modern agrosystems (*31-35*), pathogen populations are generalist and circulate extensively in the agrosystem. Such conditions of epidemiology and farming practices in traditional agrosystems would not be favorable to the coevolutionary dynamics of the “arms race” type (*36-39*), and this would explain why landraces do not present signatures of strong positive directional selection on immunity genes, unlike modern varieties.

Our study highlights the importance of preserving and studying landraces and underscores the need for targeted efforts to integrate the standing variation in landraces into new varieties. The Yuanyang terraces also provide a model from which we could draw inspiration to deploy such varieties to reduce the risk of emergence of specialized and virulent pathogenic lineages. Such endeavors hold promise for fostering sustainable agriculture in the face of evolving challenges posed by changing environmental conditions and the emergence of new plant diseases.

## Materials and Methods

### DNA sample preparation and sequencing

The 353 landrace accessions and the 42 modern varieties sequenced in this study (Dataset S1) were grown at Yunnan Agricultural University, and Plant Health Institute Montpellier, respectively. Young leaves were collected from the plants and snap-frozen in liquid nitrogen. Total DNA was extracted with the DNAsecure plant kit (TIANGEN, Beijing). 2 µg genomic DNA from each accession was used to construct a sequencing library following the manufacturer’s instructions using NEBNext Ultra DNA Library Prep Kit (NEB Inc., America). Paired-end sequencing libraries with an insert size of approximately 400 bp were sequenced on an Illumina HiSeq 4000 sequencer. 437 genomes of 3,000 Rice Genomes Project (*40*) were randomly sampled and analyzed jointly with 395 new genomes produced in this study. Paired-end resequencing reads were cleaned using fastp (Version: 0.12.2) (*41*).

### Variation calling and annotation

Paired-end resequencing reads were mapped to an *indica* rice genome Shuhui498 (R498) reference genome(*17*) with BWA (Version: 0.7.10-r789; default settings). Duplicated reads were marked with the Picard package (picard.sourceforge.net, Version: 2.1.1). Reads around indels were realigned using the Genome Analysis Toolkit (GATK, Version: 3.3-0-g37228af). Variant detection followed the best practice workflow recommended by GATK (*42*). Variants were called for each accession using the GATK HaplotypeCaller (*42*). Joint genotyping was performed on the gVCF files with parameters “QD < 5.0 || MQ < 40.0 || FS > 60.0 || SOR > 3.0 || MQRankSum < -5.0 || ReadPosRankSum < -5.0 || QUAL < 30”, and the Indel filter expression was set as “QD < 5.0 || ReadPosRankSum < -5.0 || InbreedingCoeff < -0.8 || FS > 100.0 || SOR > 5.0 || QUAL < 30”. Only insertions and deletions shorter than or equal to 40 bp were considered.

Two subsets of the SNPs were defined using the following filtering criteria: (1) a base SNP set of 17,295,637 SNPs created from the high-quality bi-allelic SNPs by removing SNPs in which heterozygosity exceeds Hardy–Weinberg expectation for a partially inbred species, using the same approach as previously described (*4*); (2) a filtered SNP set of 4,579,513 SNPs created from the ∼ 17-million-SNP base SNP set by removing SNPs with >20% missing calls and MAF < 1%. Heterozygosity was computed using Vcftools 0.1.16 (*43*). SNPs and Indels were annotated using package ANNOVAR 2015-12-14 (*44*).

Immunity-related genes were identified using InterPro annotations from R498 genome annotation V3: NOD-Like Receptors (NLRs), Wall-Associated Kinases (WAKs), and Receptor-like kinases (RLKs) other than WAKs (Dataset S5). The resistance genes were identified in the reference genome by similarity analysis with their sequences in the Nipponbare genome (https://rice.uga.edu/; Table S6), using cutoffs of 0.0001 for the E-value, 500 for the score, and 90 for the percentage identity.

### Summary statistics of genomic variation

Nucleotide diversity in 10kb windows and genes was estimated using the scikit-allel 1.3.3 Python package (*45*). Nucleotide diversity in the group of indica YYT landraces and clusters of indica YYT landraces were computed by randomly sampling one accession per landrace. Nucleotide divergence between clusters (dxy) was computed using the same subsampling approach, with scikit-allel.

### Inference of population structure

Principal component analysis was performed using the scikit-allel in Python on the filtered SNP set, with prior pruning of linked SNPs with Plink 2.0 alpha (*46*) (window size: 50 SNPs, step size: 10 SNPs, r^2^ threshold: 0.1). Population subdivision was inferred using Admixture 1.3.0 on the filtered SNP set. Admixture was run with the number of clusters K ranging from 1 to 30, and 10 repeats for each K. The q-matrices (q_i_ being the membership proportions in cluster i) inferred by Admixture were aligned and clustered based on similarity using Clumpak (*47*). The q-matrices belonging to the largest mode of each K value were averaged to produce the final matrix of membership proportions. Cluster membership for each accession was defined by applying a threshold of ≥ 0.65 to the averaged q-matrix estimated for the most biologically relevant K value (*4*): accessions were assigned to the cluster in which their q-value was the highest and ≥0.65. Accessions with q values <0.65 were classified as follows: (i) if the sum of q-values in YYT clusters was ≥0.65, the accession was classified as “YYT admixed”, (ii) if the sum of q-values in modern indica clusters was ≥0.65, the accession was classified as “XI admixed”, (iii) if both the sum of q-values in modern indica clusters and in the japonica cluster was ≥0.325, the accession was classified as “XI/JAP admixed”, (iv) if both the sum of q-values in modern indica clusters and in YYT cluster was ≥0.325, the accession was classified as “XI/YYT admixed”, (v) if both the sum of q-values in YYT clusters and in YYT cluster was ≥0.325, the accession was classified as “YYT/JAP admixed”, (vi) the remaining accessions were classified as “admixed”. Branches of the neighbor-joining tree were colored according to this classification.

### Inference of demographic history

Historical changes in effective population size were estimated using the coalescent approach implemented in MSMC2 (*19*), with a mutation rate of 6.5 × 10^-9^ /bp/generation (*20*) and a generation time of one year. For each cluster, ten independent analyses were conducted with ten random sets of eight haplotypes, each representing a different accession. In cross-cluster analyses, ten independent analyses were conducted with ten random sets of eight haplotypes, with four haplotypes from each cluster, each haplotype representing a different accession. The time sequence modeled was 1*2+30*1+1*2.

### Tests for balancing and recent directional selection

Balancing selection was analyzed using BetaScan 2 (*26*), with a sliding window size of 1000 bp and filtering sites with folded frequency below 0.01. Ancestral alleles were identified using *O. barthii* as an outgroup. The mutation rate theta was calculated as 4*Ne*µ based on the site frequency spectrum of SNPs in putatively neutrally evolving regions of the genome, where µ is the locus neutral mutation rate and Ne is the effective population size. Putatively neutrally evolving regions were identified by filtering out genomic regions corresponding to coding sequences and repetitive elements. The divergence time between *O. barthii* and clusters was estimated using Ballet (*48*).

Genomes were scanned using Selscan 2 for signatures of recent directional selection (*27*), using physical distance along chromosomes (as opposed to map distance). Prior to running Selscan, genotypes were phased using Beagle 5.4 (*49*) with default settings. Analyses were conducted with pairs including one YYT cluster and one modern indica cluster, and we calculated the cross-population extension of the haplotype-based statistic nSL (XP-NSL) (*50*).

## Acknowledgments

We are thankful to Dr. Yongsheng Liu, School of Horticulture, Anhui Agricultural University for help in collecting biological materials.

## Author contributions

Conceptualization: HH, JBM, YD Methodology: HH, JBM

Data curation: PG, SD, HH Formal analysis: PG, SD, JBM

Investigation: HH, XL, LM, XH, YZ, CL Resources: HH, JBM, YD

Visualization: SD, PG Supervision: HH, PG, JBM

Writing—original draft: PG, SD, JBM

Writing—review & editing: HH, PG, SD, JBM

Project administration: JBM, HH

Funding acquisition: HH, YD, JBM

## Data availability

Raw reads are available at the China National Center for Bioinformation under the accession number PRJCA030930. VCF files listing SNPs and the results of genome scans for balancing and directional selection are openly available at http://teabase.ynau.edu.cn/data/tea/pangeome/rice/

## Supporting information

**Dataset S1**.

Sample information.

**Dataset S2**.

Proportions of ancestry of 832 Asian rice accession in K=2 to K=30 clusters, as estimated using the Admixture program. HighestGcluster is the cluster in which a given accession had highest membership at K=10. Hybrid/admixed identifies accessions that are Japonica-indica hybrids, or have membership in multiple Indica lineages.

**Dataset S3**.

RLK, NLR, WAK, Resistance and other genes in the top 5% of ß-scores.

**Dataset S4**.

RLK, NLR, WAK, Resistance and other genes in the top 5% of XP-NSL scores.

**Dataset S5**.

NOD-Like Receptors (NLRs), Wall-Associated Kinases (WAKs), Receptor-like kinases (RLKs) other than WAKs, and resistance genes (R-genes)

**Figure S1**.

Cross-validation error and frequency of major mode as a function of the number of ancestral populations K, as estimated using the clustering algorithm implemented in Admixture. For each value of K, the major mode represents the main clustering solutions among 30 replicates.

**Figure S2**.

Population subdivision in 832 Asian rice accessions, as inferred using the program Admixture, with the number of modeled clusters ranging from K=2 to K=30. Each vertical bar represents an individual accession, with colors indicating the proportion of ancestry from each of the K inferred clusters.

**Table S1**.

Average absolute divergence (dxy) per base pair between Asian rice accessions from different clusters

**Table S2**.

Nucleotide diversity per base pair estimated in 10kb windows. Each landrace was represented by a single, randomly sampled, accession.

**Table S3**.

Nucleotide diversity per base pair estimated in genes. Each landrace was represented by a single, randomly sampled, accession.

**Table S4**.

Number of RLK, NLR, WAK and Resistance genes in the top 5% of beta scores in clusters of modern indica and indica landraces.

**Table S5**.

Number of RLK, NLR, WAK and Resistance genes in the top 5% of XP-NSL scores. Analyses were carried out on pairs of clusters and thus counts are reported for each pair of clusters (“contrasts”). Counts per cluster were obtained by counting genes in the top 5% of XP-NSL in all contrasts in which the cluster was included.

**Table S6**.

Identifiers of R-genes in Nipponbare (https://rice.uga.edu/) and R498 assemblies.

