## Supplementary material for "Genome Sequencing of Rice Landraces from the Yuanyang Terraces Uncovers Ancient and Diverse Lineages of Indica Rice": Figure S2

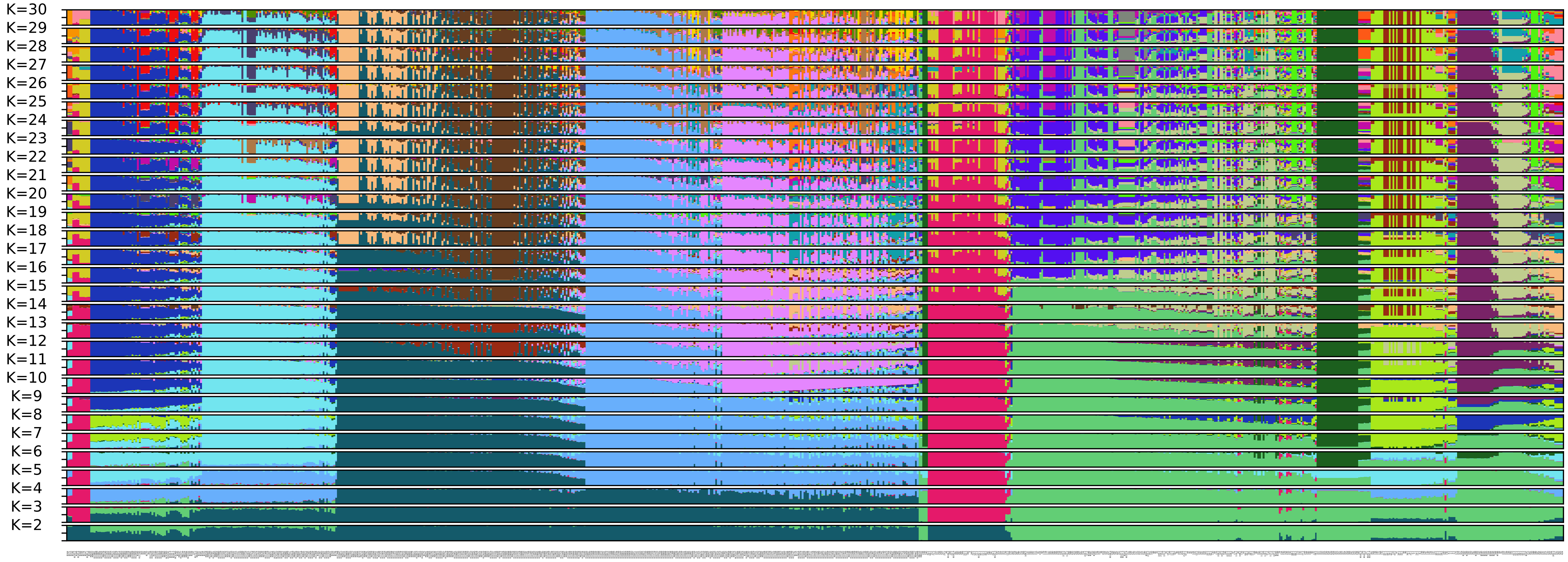


Figure S2: Population subdivision in 832 Asian rice accessions, as inferred using the program Admixture, with the number of modeled clusters ranging from K=2 to K=30. Each vertical bar represents an individual accession, with colors indicating the proportion of ancestry from each of the K inferred clusters.
