## Supplementary material for "Genome Sequencing of Rice Landraces from the Yuanyang Terraces Uncovers Ancient and Diverse Lineages of Indica Rice": Figure S1

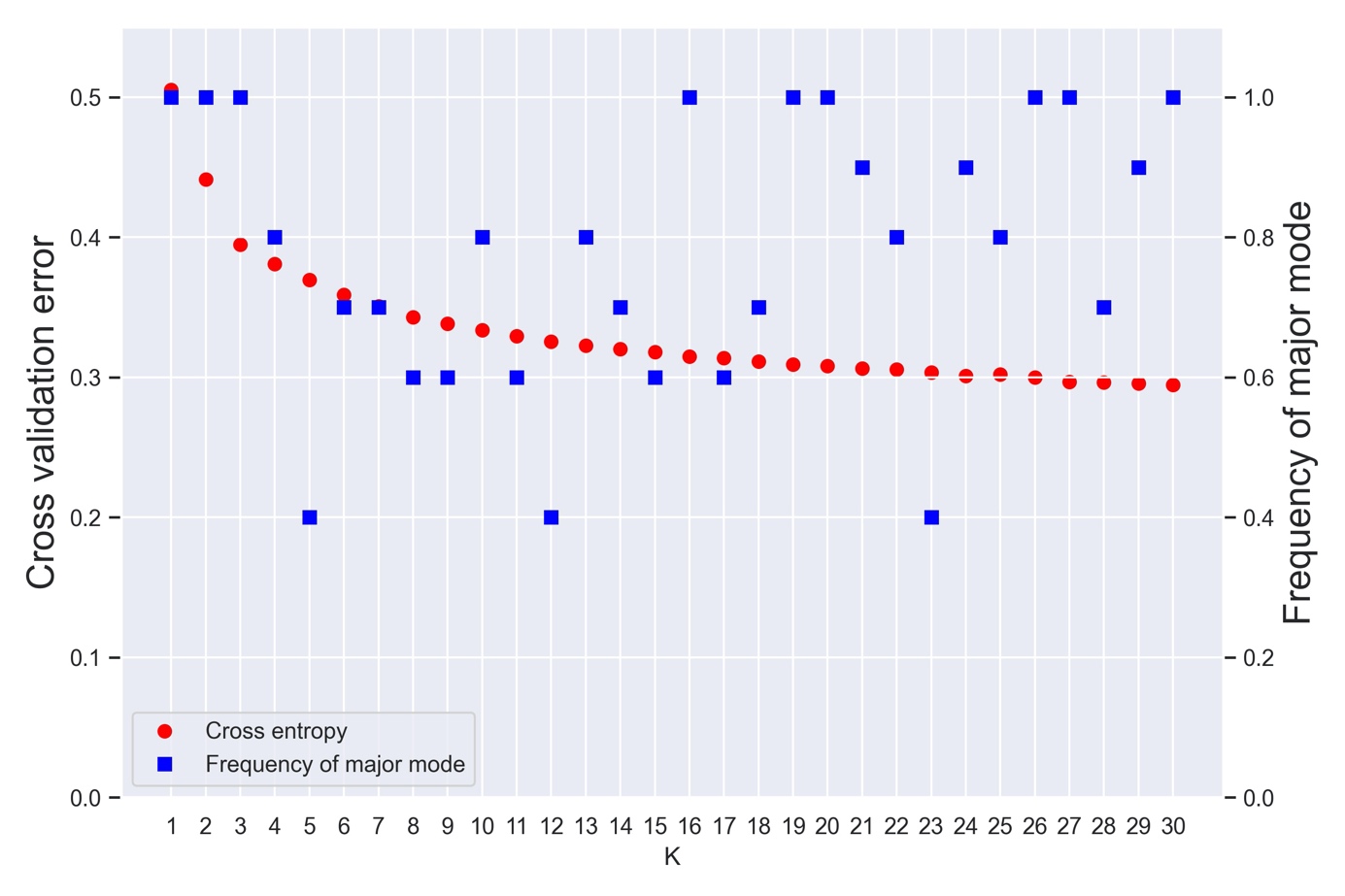

Figure S1. Cross-validation error and frequency of major mode as a function of the number of ancestral populations K, as estimated using the clustering algorithm implemented in Admixture. For each value of K, the major mode represents the main clustering solutions among 30 replicates.
